# Asymmetric division and polar growth define a non-canonical cell cycle in *Bradyrhizobium*

**DOI:** 10.64898/2026.11.05.764102

**Authors:** Ian F. Medici, Julieta Bressano, Mounia Kortebi, Diego J. Comerci, Elías Mongiardini, Virginia S. Lioy, Alfonso Soler-Bistué

## Abstract

Bradyrhizobia are widely used as plant growth promoters in agriculture, yet their cell biology remains poorly understood. Here, we describe, for the first time, nucleoid segregation dynamics and the cell cycle of *Bradyrhizobium* at the cellular level. We show that highly asymmetric polar growth generates a short daughter cell and a longer mother cell at each division. Nucleoid compaction and positioning change across growth phases, culminating in a highly compacted, asymmetrically positioned nucleoid in stationary phase. ParB presents a subpolar localization, at the nucleoid periphery. Despite this pronounced cellular asymmetry and the presence of a single large chromosome, chromosome conformation capture reveals a global chromosome organization similar to that of bacteria undergoing conventional binary fission. Together, these findings uncover a distinctive cell cycle program in *Bradyrhizobium*, showing that such extreme asymmetry in growth and division, significantly different from that observed in closely related characterized α-Proteobacteriota, is compatible with conserved principles of chromosome organization.

## Introduction

Bacteria have been studied for decades to understand the coordination between cell growth and division[1–3]. Much of this knowledge derives from a few fast-growing useful laboratory models, particularly *Escherichia coli*, *Bacillus subtilis* and *Caulobacter crescentus*[1, 4, 5]. However, they represent only a narrow fraction of the full bacterial diversity[6, 7], which spans an extraordinary range of ecological niches and growth strategies[8–11], suggesting that alternative cell-cycle programs remain unexplored. Advances in flow cytometry, microscopy and microfluidics have been critical for gaining insight into the molecular mechanisms defining cell growth and division[12–14]. Nevertheless, slow-growing microorganisms remain particularly difficult to analyze, and little is known about how they coordinate growth, replication and division at the single-cell level[15].

The dynamics of bacterial chromosome organization have also been studied throughout the cell cycle in bacteria[16–21]. Bacterial chromosomes adopt a distinct three-dimensional (3D) architecture, with individual chromosomal loci occupying defined intracellular positions (*i.e.*, specific chromosome configurations) and following a highly coordinated choreography during chromosome segregation and cell division. This process is orchestrated by protein complexes that contribute to chromosome segregation, among which the ParABS system is particularly notable and broadly conserved in bacteria. In most of them, it positions the *origin* region (*ori*) and drives its faithful segregation[22–27]. Frequently, the presence of the ParABS system is correlated with chromosomes with a longitudinal disposition inside the cell (the origin and terminus regions of the chromosome are positioned at opposite poles) and with the presence of a specific 3D architecture in which long-range DNA contacts occur both in *cis* (on the same arm of the chromosome) and in *trans* (between the two chromosome arms)[17, 28–31]. Contacts *in trans* are usually mediated by the Smc-ScpAB complex, which is specifically loaded near the origin of replication by the ParB/*parS* complex[17, 29, 31–34].

Bradyrhizobia are arguably the most abundant microorganisms in soils [35]. They are critical in agriculture due to their plant growth-promoting capabilities in important crops such as peanut and soybean[36–38]. With doubling times ranging from 9 to 40 hours, they are among the slowest-growing culturable bacteria[39]. This trait complicates their study and their genetic modification. As a result, *Bradyrhizobium* physiology remains poorly explored and, at the cellular level, there are almost no studies done[40, 41]. *Bradyrhizobium* belongs to α-Proteobacteriota, which include the well characterized model bacterium *C. crescentus,* which exemplifies cell asymmetry, and has a particular cell cycle associated with a two-stage lifestyle[4]. *C. crescentus* mother cells are attached to a surface, releasing structurally different daughter cells displaying a flagellum. Newly released cells block DNA replication at the G1 phase while they explore the environment to find an appropriate niche. Even if *Bradyrhizobium* and *C. crescentus* are distantly related within the α-Proteobacteriota, key regulators of the cell cycle are highly conserved[42]. Within this latter group, several bacterial species belonging to *Hyphomicrobiales* (formerly known as *Rhizobiables*) such as those from the *Agrobacterium, Sinorhizobium,* and *Brucella* genera share remarkable cell-cycle properties such as the asymmetric position of the septation site and differential presence of proteins according to bacterial old or new poles[43]. Many aspects of cell division and chromosome segregation are species-specific properties and remain to be fully characterized[44]. For example, most *Hyphomicrobiales* harbor multiple chromosomes, a trait shared with 10% of bacteria and particularly frequent in those interacting with a host[45, 46]. We recently reported that *Bradyrhizobium,* contrary to classical models, reduces its size in the transition from lag to exponential phase; then cells increase their length again in stationary phase[40], raising the question of how chromosome segregation and organization are orchestrated during these cellular gymnastics. Here, we characterize *Bradyrhizobium japonicum* E109 (Bj109) chromosome organization, segregation and cell division, providing a comprehensive view of the cell cycle in this challenging-to-handle, and biotechnologically relevant model. Overall, we unveil that Bj109 presents an extreme cell growth polarity coupled with a marked asymmetric division cycle that differs from those described in other α-Proteobacteria.

## Results

### *B. japonicum* presents an extreme polar growth with asymmetrical cell division

To characterize nucleoid segregation dynamics in Bj109, we grew cells up to early exponential phase (EEP), exponential phase (EP) and early stationary phase (ESP) and then we fixed them and stained the nucleoids with Hoechst 33342 (Figure 1A). The quantitative analysis and statistical comparison of differences in cell size and nucleoid compaction along EEP, EP and ESP were then performed (Figures 1B, 1C and 1D respectively). We found that Bj109 cell length varies from 2.8 ± 0.9 μm during EEP to 2.1 ± 0.6 μm in EP, finally reaching 2.4 ± 0.9 μm in ESP (Figures 1B and 1C, Table S2). Analysis of Hoechst 33342 fluorescence intensity across cell lengths revealed a drastic nucleoid compaction along the growth stages (Figure 1B and 1D, Table S2). Quantification of Hoechst 33342 intensity revealed that the signal occupied 50.8 ± 1.3%, 38.6 ± 1.3 % and 32.6 ± 1.3% of the cell area during EEP, EP, and ESP, respectively (Figure 1D. Table S2). Furthermore, DNA compaction is correlated with an asymmetric localization of the nucleoid within the cell, especially in ESP (Figure 1B and Table S2). This pattern does not depend on the growth medium since we observe similar behavior in cells grown in YEM medium (Figure S1. Table S3). Overall, we confirmed that Bj109 cell size changes throughout the cell cycle[40] and revealed a gradual increase in nucleoid compaction as the cell cycle progresses.

**Figure 1:**
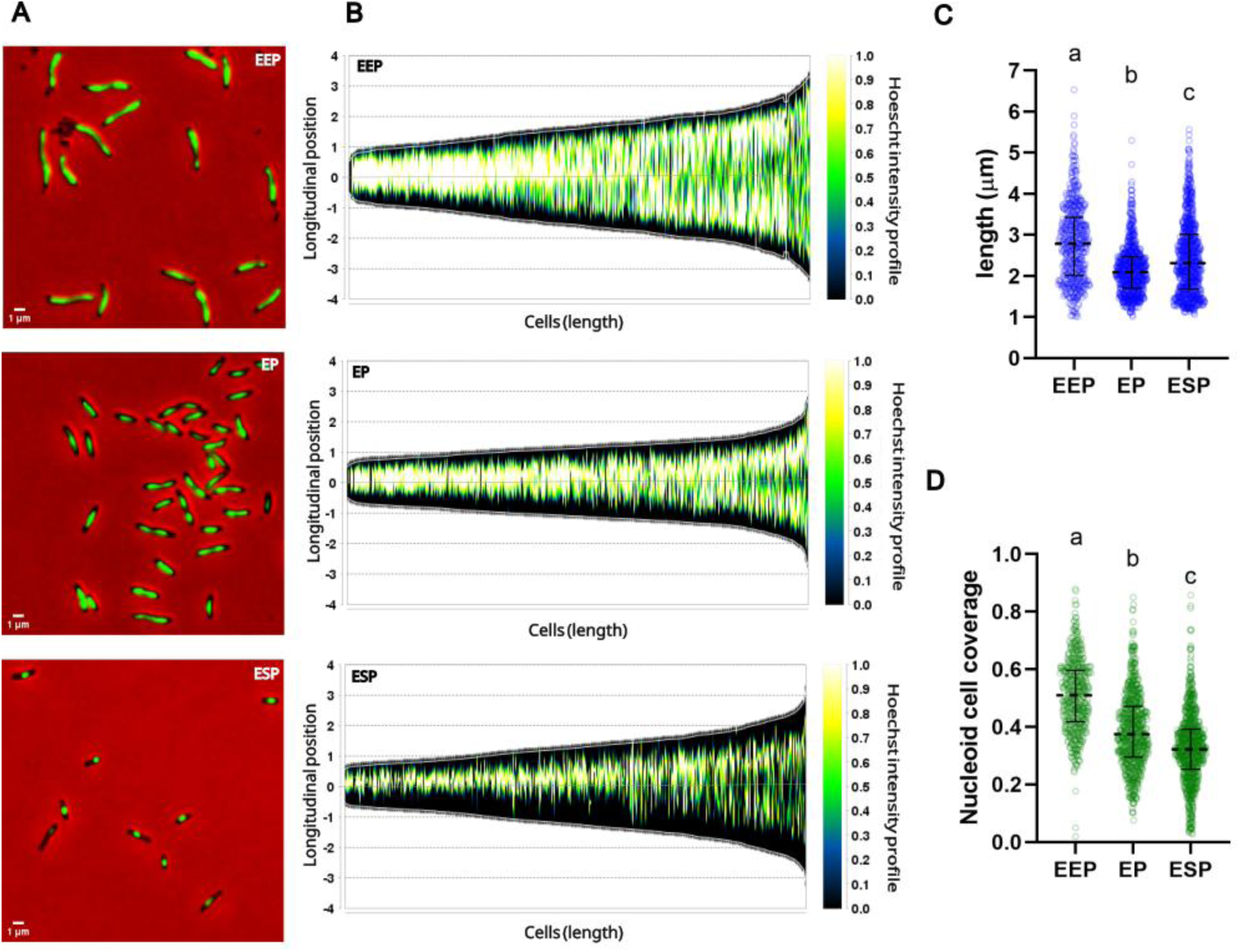
Cell size and nucleoid distribution vary along the growth phases of Bj109. **A.** Representative image of cells stained with Hoechst 33342 at different stages of growth in AG medium. **B.** Demographic representation showing the Hoechst 33342 intensity profiles of cells sorted by length at the indicated phases. **C.** Distribution of cell sizes at the indicated phases. **D.** Proportion of the cell covered by Hoechst 33342 intensity at the indicated phases. The middle lines indicate the median for each timepoint. Bars show the quartile range. Letters indicate statistically different groups (Kruskal-Wallis and Dunn’s tests, p<0,0001). For panels **B** to **D** the number of analyzed cells is 547 for EEP, 759 for EP and 861 for ESP.

To further characterize Bj109 cell division throughout the cell cycle, we grew cells until EP and then exposed them to a pulse of fluorescent D-amino acids (FDAA)[47, 48], allowing the labeling of the newly synthesized cell wall. Cells were subsequently fixed, and stained with Hoechst 33342, enabling the simultaneous visualization of nucleoid segregation and cell wall synthesis at the single-cell level (Figure 2A-B). Analysis of labeled cells unveils that cell wall synthesis is highly asymmetric, particularly in larger cells (longer than 3.5 µm) containing two well-defined nucleoids, a hallmark of the stage immediately preceding cell division (Figure 2C-D). To further explore FDAA incorporation and its link with nucleoid segregation, we plotted the density distribution of FDAA maxima and nucleoid location, as a function of the relative longitudinal axes of cells bearing either one or two nucleoids (Figure 2E). This analysis revealed distinct patterns of cell wall synthesis during the cell cycle. Cells containing a single nucleoid show a higher FDAA fluorescence density at one pole while the opposite pole presents a less dense signal. Additionally, in single-nucleoid cells, nucleoid location density is asymmetrically positioned with its maximal density offset from mid-cell. In contrast, large cells containing two nucleoids, corresponding to the pre-divisional stage, display a major density of FDAA signal at one pole, together with a second site of incorporation at the developing septum. Notably, this septal signal is shifted from mid-cell, indicating that septation occurs asymmetrically (Figure 2A-2E). Consistent with this pattern, the density distribution of nucleoid locations in cells bearing two nucleoids was also asymmetric relative to midcell.

**Figure 2:**
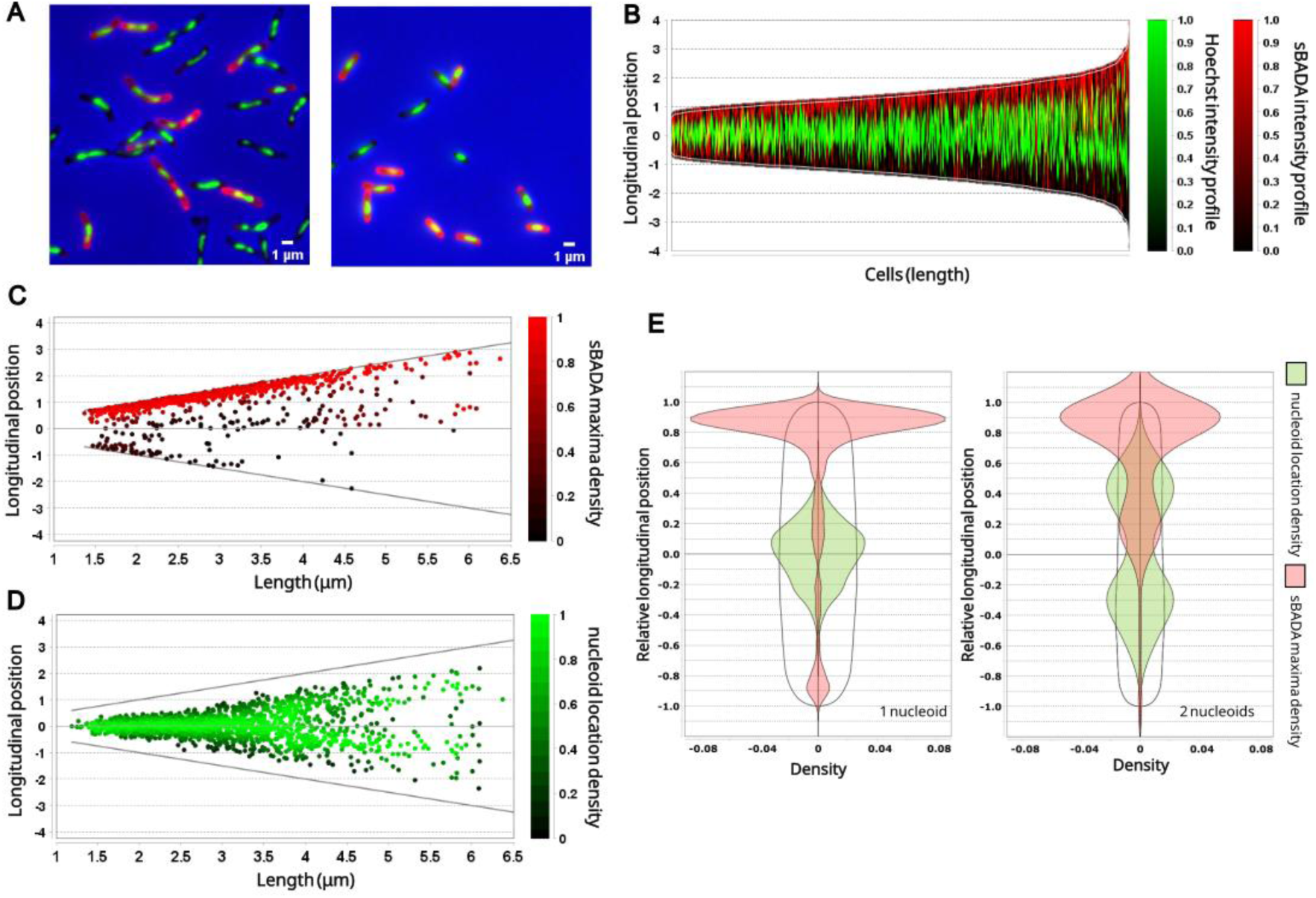
Cells display extreme polar growth and asymmetrical division. **A.** Representative fluorescence microscopy image of cells in EP grown in AG medium in EP. Cells were stained with the FDAA sBADA to detect cell wall synthesis, and with Hoechst 33342 for DNA. **B.** Demographic representation showing the sBADA and Hoechst 33342 intensity profiles of cells sorted by length and oriented by sBADA maximal intensity (N = 1427). **C.** Distribution and density of sBADA maxima within the cells sorted by length. **D.** Distribution and density of nucleoid location within cells sorted by length. **E.** Cell density distribution plot of sBADA maxima (red) and nucleoid location (green) as a function of relative longitudinal axes of cells grouped by the number of nucleoids detected. Straight lines at coordinate zero of the vertical axes correspond to midcell. For panels **B** to **E** cells were oriented according to sBADA signal maximum intensity.

To further characterize Bj109 asymmetric cell division and to distinguish cell elongation from septation, we sequentially labeled cells with two different FDAAs. Microscopy images show that HADA, the first stain, accumulates on one half of the cell, while sBADA, the second stain, accumulates at the tip of the same half of the cell and in the developing septum (Figure 3A). To quantitatively unveil the dynamics of cell elongation and septation, we plotted the density of HADA and sBADA maxima within the cell, as a function of cell length (Figure 3B & C). In agreement with unipolar cell enlargement, in small cells (shorter than 3.5 µm), the HADA signal is subpolar compared to sBADA fluorescence. Larger cells (longer than 3.5 µm) display a similar distribution of HADA and sBADA at the growing pole, and a second site of sBADA incorporation at the developing septum. Similar results were obtained when inverting the order of the FDAA incorporation (Figure S2).

**Figure 3:**
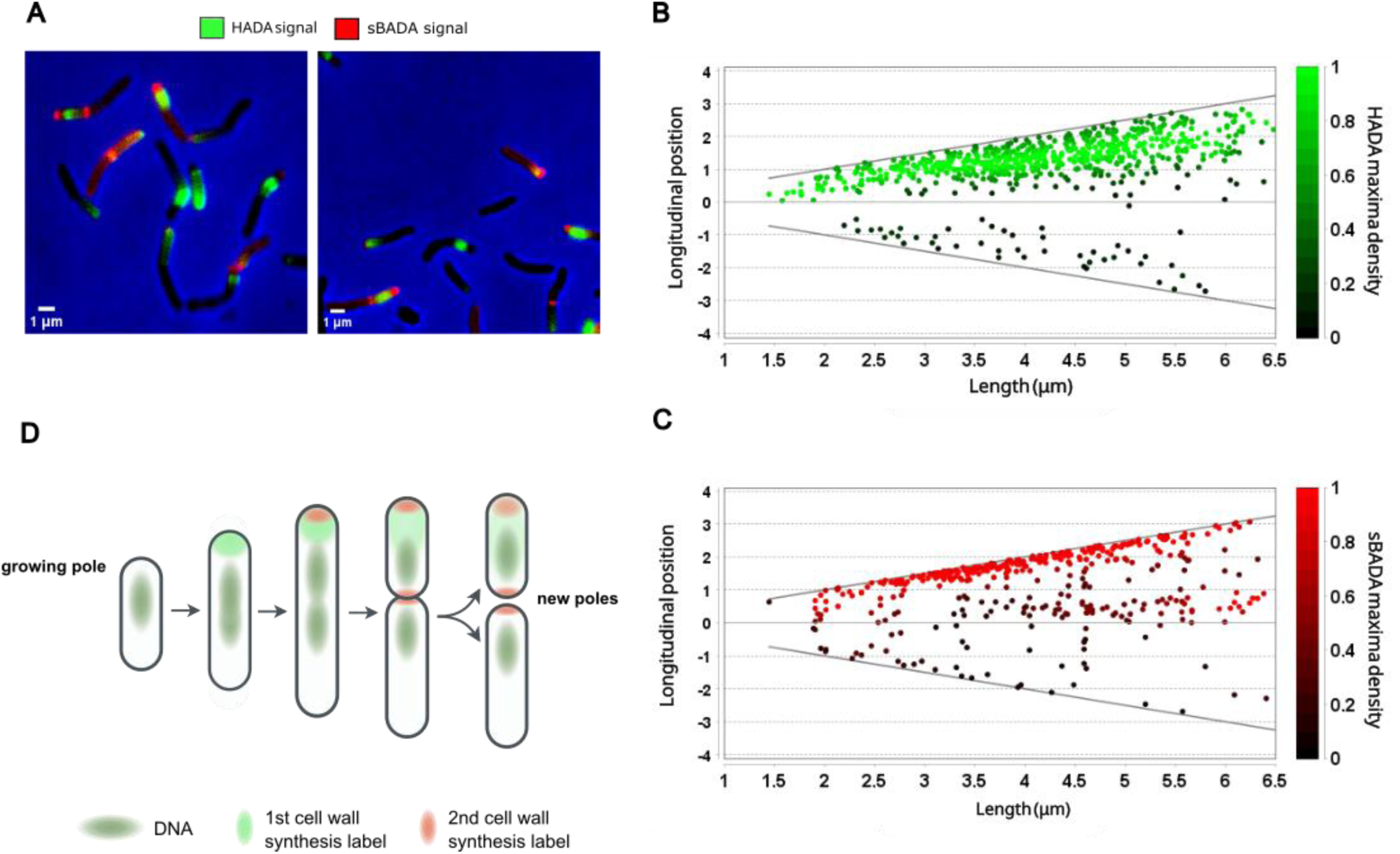
Cell wall synthesis occurs in two defined and sequential steps. **A.** Representative fluorescence microscopy image of Bj109 cells during EP in AG medium. Cells were sequentially stained with the FDAAs HADA and then sBADA. **B & C.** Distribution and density of HADA and sBADA maxima, within the cells sorted by length and oriented by the maximum intensity of FDAA signal (N = 1152). **D.** Schematic representation of cell growth, division and DNA segregation hypothesized from results.

Overall, these results show that cell elongation occurs exclusively at one pole - the growing pole - and is followed by peptidoglycan synthesis at the septum, which gives rise to a new pole upon division. In descendant cells, the nucleoid is asymmetrically positioned, slightly closer to the new pole, presumably because septation occurs shortly after chromosome segregation. Asymmetric division at approximately two-thirds of the cell length produces a large mother cell and a small daughter cell, resulting in descendant cells with distinct sizes and polar histories. The large cell inherits one pre-existing pole and one newly formed pole generated during septation, whereas the small cell inherits both the new pole generated during septation and the growing pole that underwent cell wall synthesis during the preceding elongation phase. Consequently, the two daughter cells may resume the cell cycle from different initial configurations, contributing to the heterogeneity of the exponentially growing population[41].

### ParB segregation correlates with polar growth and precedes septum synthesis

To uncover how chromosome segregation is orchestrated in a bacterium displaying such an extremely asymmetric mode of cell division, we investigated the subcellular localization of ParB. ParB specifically binds to *parS* sequences, which are typically located close to the origin of replication in bacteria[27]. Together with ParA, the ParB/*parS* complex is responsible for chromosome segregation in most bacterial species[22]. Following DNA replication, the duplicated ParB/*parS* complexes are rapidly segregated into daughter cells. In Bj109, in silico analyses using the 5′-TGTTNCACGTGAAACA-3′ consensus[27] revealed 2 *parS* sites located at positions 624,137 and 624,152 bp, respectively, within the *oriC* region (624,080 to 624,665 bp). Therefore, ParB localization can be used as a proxy for origin segregation. To this end, we constructed a merodiploid strain carrying a second copy of the *parAB* operon (RN69_RS02875 and RN69_RS02880 genes, respectively, see Figure S3 and Supplementary Text), in which *parB* was fused to the gene encoding yellow fluorescent protein (*yfp*) (Figure S3). This strain, hereafter referred to as YFP-ParB, displayed growth comparable to that of the wild-type strain with a minor increase in the lag phase duration (Figure S4). YFP-ParB localization was monitored in exponentially growing cells together with cell wall synthesis using a short HADA pulse (Figure 4A).

**Figure 4:**
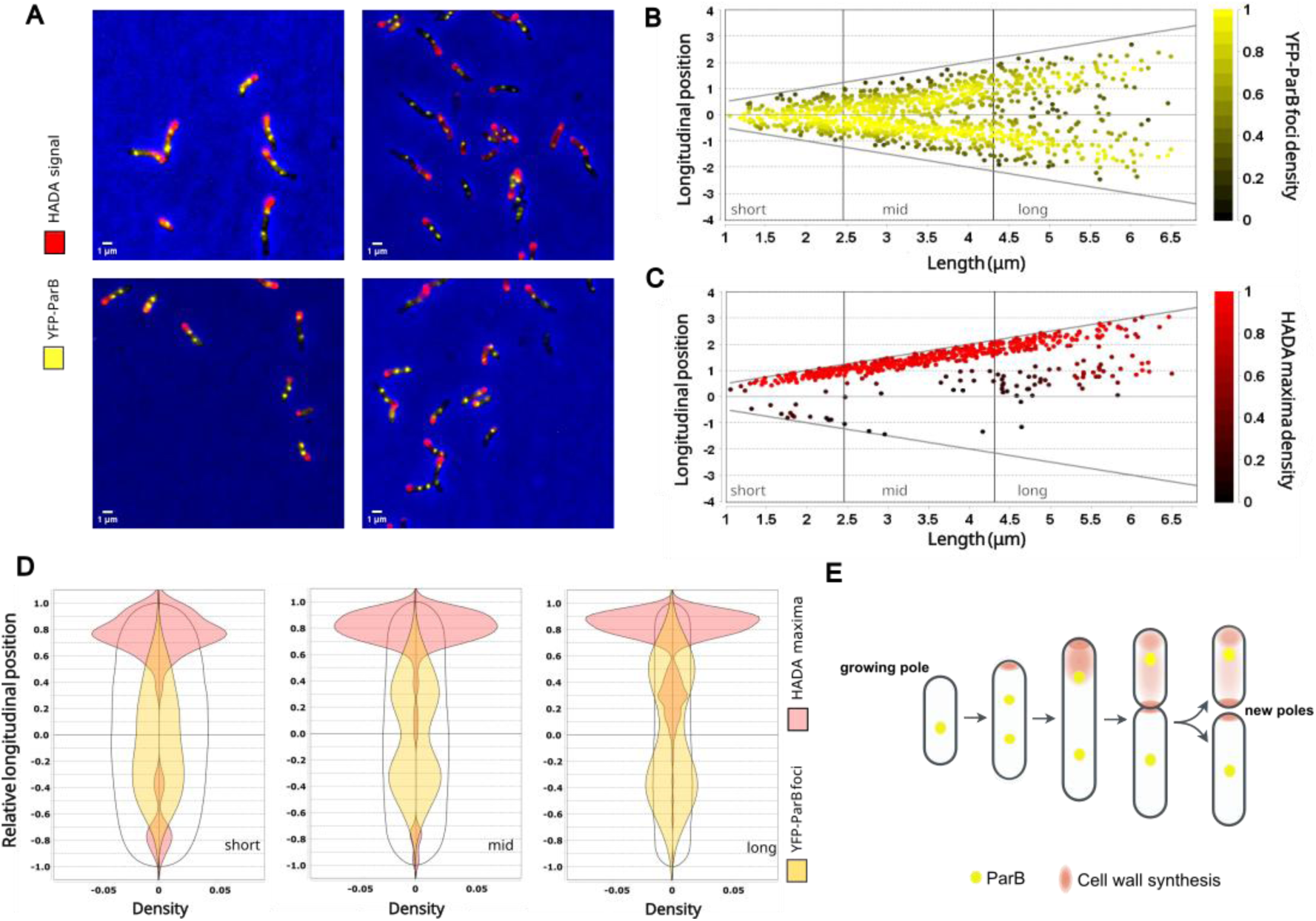
ParB segregation correlates with polar growth. **A.** Representative fluorescence microscopy image of cells during EP in AG medium. Cells expressing YFP-ParB, were stained with the FDAA HADA to detect cell wall synthesis and orient them by the labeled pole. **B.** Distribution and density of YFP-ParB foci within the cells sorted by length. **C.** Distribution and density of HADA maxima within the cells sorted by length. **D.** Cell density distribution plot of HADA maxima (red) and YFP-ParB foci (yellow) as a function of relative longitudinal axes of the cells. Cells were classified by length as shown in B & C. The total number of cells analyzed was 715, were 60% of total cells analyzed had 2 foci and of these cells 72% were long, 25% mid and 3% short **E.** Schematic representation of cell growth, division and DNA segregation according to the conclusions drawn from image analysis. For panels **B** to **D** cells were oriented according to HADA signal maximum intensity.

To quantitatively characterize ParB segregation, we plotted the densities of the YFP-ParB foci and the HADA signal maxima within the cell, as a function of cell length (Figures 4B and 4C). Cells were grouped according to their size, and the maxima density distributions of the YFP-ParB and HADA signals were plotted as a function of their relative longitudinal axes (Figure 4D). Short, intermediate-length, and long cells were defined as those below the first quartile (Q1), between the first and third quartiles (Q1– Q3), and above the third quartile (Q3) of the cell length distribution, respectively. In short cells, the YFP-ParB focus was localized along the longitudinal axis of the cell with a bias towards the side opposite to their growing pole (Figure 4B–4D). Cells of intermediate length showed a progressive accumulation of the second YFP-ParB focus towards the growing pole (Figure 4B–4D). Finally, in the longest cells, which had already initiated septum formation, two YFP-ParB foci were detected, one on each side of the developing septum (Figure 4B–4D). Analyses of YFP-ParB position after Hoechst 33342 staining (Figure S5), reveal that in all cells, YFP-ParB foci are located in regions with low DNA intensity and close to the poles (Figure S5B-5D). However, YFP-ParB foci show limited colocalization with HADA labeling, indicating a subpolar localization (Figure 4A-C). Furthermore, in cells with two foci, the distance between them increases together with cell size and synthesis of the septum. All in all, these results suggest that after replication, the ParB/*parS* complex is segregated towards the future daughter cells concomitantly with polar growth, and once the ParB/*parS* complex has been segregated, septum synthesis is initiated (Figures 4E and S5E).

To complement our analysis of ParB segregation, we next examined the global organization of the chromosome in exponentially growing Bj109 cells using high-resolution chromosome conformation capture (Hi-C). Contact maps generated at 5-kb resolution revealed a main diagonal representing contacts between neighboring genomic loci and a secondary diagonal corresponding to interactions between the two chromosome arms (Figure 5A). Thus, in addition to the coordinated positioning of the origin region during polar growth and division, Hi-C revealed inter-arm contacts as a prominent feature of chromosome organization under these growth conditions. Altogether, these results reveal that Bj109, despite its slow replication rate, marked asymmetric cell division and large circular chromosome (>9 Mb), presents a large-scale chromosome organization comparable to that observed in bacteria with smaller circular chromosomes, binary fission and faster replication rates.

**Figure 5.**
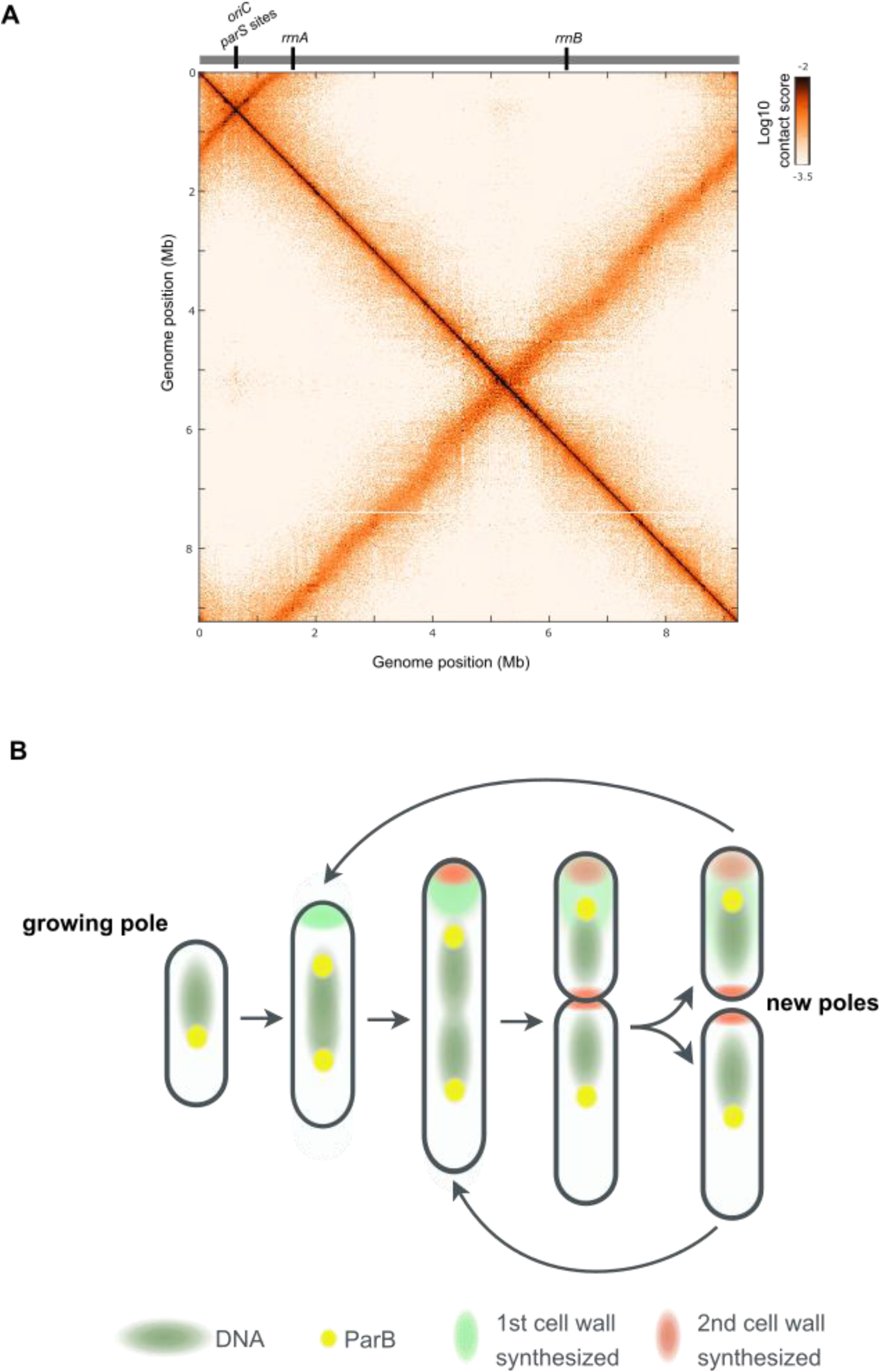
3D chromosome organization and a model for asymmetric cell division during the Bj109 cell cycle. **A.** Genome-wide Hi-C contact matrix at 5-kb resolution for Bj109 cells growing in EP. The main diagonal reflects contacts between adjacent genomic loci, whereas the secondary diagonal corresponds to interactions between loci on opposite chromosome arms. Contact frequencies are shown as log-transformed normalized values, where white represents low contact frequencies, orange intermediate frequencies, and black the highest interaction frequencies. The positions of *oriC*, the *parS* sites, and the rDNA operons are highlighted above the matrix. **B.** Model of unipolar growth and chromosome segregation during the Bj109 cell cycle. Newborn cells contain a single compact chromosome (green shading) and one YFP-ParB focus (yellow). As the chromosome replicates, the two YFP-ParB foci separate, with one focus moving in concert with the growing pole. Asymmetric cytokinesis produces a smaller daughter cell and a larger mother cell. YFP-ParB foci localize to regions of low DNA signal, likely at the edges of the nucleoid, consistent with a model in which ParB pulls the segregating chromosome toward the growing pole. After cell division, daughter and mother cells initiate a new round of cell elongation at the new pole of each cell.

## Discussion

Knowledge of cell cycle details is critical for understanding the ecology, the physiology and the homeostasis of single cells with potential applications ranging from synthetic biology to bioremediation and cancer[49–51]. The cell cycle, and particularly the relationship between cell size, shape and DNA segregation, has been extensively studied in several model organisms in across the three domains of life, including archaea, *E. coli, B. subtilis, S. cerevisiae* and human cells[52–55]. Recent work has explored these topics in less studied models with unusual shapes, ploidy, habitats or ecological strategies displaying a great variety of potential ways for the cell cycle to proceed (e.g.*, [56–60]*)

Bj109 belongs to the α-Proteobacteriota, particularly to the order *Hyphomicrobiales*, which encompasses some well-studied model organisms such as *Agrobacterium[61, 62]*, *Sinorhizobium[63, 64]* and *Brucella[43, 65, 66]*. *Hyphomicrobiales* are microorganisms interacting with hosts. They frequently present multiple chromosomes and they are paradigms of unipolar elongation and incorporation of peptidoglycan at one end of the rod[43]. Bj109 is characterized by slow growth and a single large circular chromosome of 9.22 Mb[67]. Despite its ecological and economic importance, its cell cycle remains poorly characterized, partly because of its extremely slow growth difficulting experimental manipulation and genetic modification. Here, we provide the first detailed description of the Bj109 cell cycle. While apparently respecting the G1-S-G2 cell cycle observed in other alphas[44, 68], Bj109 displays unique characteristics. For instance, the asymmetry at cell division seems to be more pronounced than in other *Hyphomicrobiales.* Additionally, cell size variation changes drastically along growth phases, showing a mother cell much longer than newborn cells (Figures 1 and 2). The synthesis of the cell wall occurs sequentially, first almost exclusively at the growing pole and in a second step at the developing septum (Figure 3).

We also studied Bj109 chromosome segregation along different growth phases, uncovering a progressive increase in chromosome compaction along the cell cycle (Figure 1). Furthermore, in early stationary phase, the chromosome of Bj109 was not only compacted but also asymmetrically localized, particularly in small cells. To further understand nucleoid segregation, we investigated the localization of YFP-ParB positioning during exponential phase, unveiling a particular segregation dynamics.

Altogether, these results support a model (Figure 5B) in which cells are born with a single YFP-ParB focus and a compacted chromosome. Unipolar cell elongation is accompanied by chromosome replication and YFP-ParB focus duplication and segregation, followed by asymmetric septation that gives rise to a large mother cell and a small daughter cell. This division choreography is compatible with a process in which chromosome replication initiates once per cell cycle. The nucleoid distribution in predivisional cells suggests a bias in the positioning of the nucleoid destined for the smaller daughter cell towards the developing septum (future new pole), whereas in cells containing a single nucleoid, its position is shifted towards the elongating pole (Figure 2). In parallel, the FDAA labeling pattern reveals a strong asymmetry in cell wall synthesis, with labeling predominantly restricted to one pole across the population (Figure 2-4). Notably, the occasional labeling of both poles is mainly observed in smaller cells, consistent with the simultaneous detection of cell wall synthesis associated with elongation at the pre-existing growing pole and septum formation at the opposite pole. Based on these observations, we hypothesize that, following division, both descendant cells resume polar elongation at their newly formed poles, which become the growing poles for the subsequent cell cycle. Importantly, asymmetric division gives rise to cells with distinct initial configurations, differing not only in size but also in the history of their poles. These differences may influence how descendant cells progress through the subsequent cycle, potentially leading to distinct growth trajectories or differences in the timing of the next division. This possibility highlights the importance of understanding how cell polarity is established and inherited, how it contributes to population heterogeneity[41], as well as its potential biological significance in the context of bradyrhizobia lifestyle. Finally, our results show that despite its marked asymmetric cell division cycle and large genome content, Bj109 exhibits 3D chromosome folding similar to that of other bacterial models with canonical cell division cycles.

In most bacteria, ParB positioning is directly linked to *parS* location, usually close to the origin of replication. Specific recognition of and binding to the *parS*-containing region by ParB in Bj109 remain to be experimentally demonstrated. Furthermore, in α-bacteria such as *A. tumefaciens[69], Sinorhizobium meliloti[70]*, *C. crescentus[70]* and *Brucella abortus[65]*, the origin region is positioned at the tips of the cell, while in Bj109 the origin region might be positioned at cell quarters. Future studies employing Fluorescent Repressor Operator Systems (FROS) will help us to reconcile such apparent differences by tracking the segregation choreography of specific chromosomal loci marking the *oriC*, *ter*, right and left arms[71].

Although the cell elongation pattern and the dynamics of chromosome segregation differ between Bj109 and other closely related bacterial models mentioned above, the genes regulating the cell cycle such as *ctrA, pleC, divK, ccrM* and *gcrA* among many others are well conserved among the clade[72]. However, their specific role at the molecular level is just starting to be explored within each specific model organism[61, 66, 73–76]. A limitation of this study is that the data presented come from fixed cells and, in this regard, approaches studying cells in vivo will be a great asset. For example, the use of nucleoid-associated proteins translationally fused to fluorescent proteins would confirm our observations and provide more insight into the Bj109 cell cycle. These approaches coupled with video microscopy would allow us to study all cell components in vivo, providing more precision to the study of the Bj109 cell cycle. Importantly, our work is limited to laboratory culture conditions that may drastically vary during stimulation by plant flavonoids, the contact with and attachment to the root and, most importantly, the replication inside the nodule as a bacteroid[77–79]. In conclusion, this work expands our understanding of the diversity of bacterial cell-cycle strategies beyond well characterized model organisms.

## Methods

### Growth conditions and quantification

The main strain used in this study is *Bradyrhizobium japonicum* E109 which was kindly provided by Dr. Fabricio Cassán. The derivative strains built from this isolate are detailed in Table S1 in the supplements. Bj109 strains were grown aerobically at 28°C in yeast extract-mannitol (YEM), or the AG medium[40]. For plate growth BjE109 strains were routinely grown on solid YEM at 28°C. Total biomass was estimated by measurement of the optical density at 450 nm (OD_450nm_). Samples were taken when OD_450nm_ was 0.1-0.15 for EEP, 0.3 - 0.6 for EP, and 2 – 2.5 for SP. The number of viable bacteria was estimated by counting CFUs on YEM agar plates supplemented with Congo Red (10 mg/L), using the microdilution method. Selection was made on YEM plates for BjE109 and LB plates for *E.coli*, supplemented with 200 and 50 µg/mL of kanamycin (Km), for Bj109 and *E.coli*, respectively, and 20 µg/mL of chloramphenicol for Bj109.

### Fluorescent D-amino acid (FDAA) labeling

The procedure was performed as described in[48]. Briefly, a 750µL aliquot of liquid culture at the indicated growth phase, was resuspended in 100 µL of fresh medium and 5 µL of an FDAA stock solution (5 µM) were added (Tocris Bioscience). As FDAA, green sulfonated BODIPY-FL 3-amino-D-alanine (sBADA) or blue HCC-amino-D-alanine (HADA) were used as indicated. This was incubated for 10% of the generation time. Cells were then washed twice in PBS, before analysis by fluorescence microscopy or subsequent DNA staining.

### DNA staining

Cells at the indicated growth phase were fixed by mixing the culture 1:1 with 4% paraformaldehyde (PFA) and incubating for at least 30 min at room temperature. Fixed cells were washed twice with PBS. Cell pellets were then resuspended in GTE buffer (45 mM glucose, 20 mM Tris-HCl pH 8.0, 10 mM EDTA pH 8.0) using a volume proportional to the cell density. After 1 µL of Hoechst 33342 (1mg/mL) was added to each sample, the samples were briefly vortexed and were incubated in the dark for 5 min. Samples were then resuspended in 1× PBS using a volume proportional to the cell density and analyzed by fluorescence microscopy.

### Generation of the YFP-ParB derivative

To generate the strain YFP-ParB-Bj109, we inserted a second copy of *parAB* with a parB translational fusion to codon adapted *yfp* using the plasmid pK18*mobsacB[80]*, as indicated in Figure S3. (See supplementary text).

#### Microscopy

Bacterial samples from at least 2 independent cultures for each label analyzed, were placed on a PBS agar 1.5-2% m/v slide and observed at the indicated growth phase. For cell size and fluorescence determinations, we employed snapshot analyses. Images of at least 500 cells were acquired using a DM6000-B (Leica) microscope coupled to a Prime BSI camera (Roper), and they were analyzed using MicrobeJ[81]. The feature detection tool was used for the analysis of FDAA maxima, YFP-ParB foci and nucleoid positions.

#### 3D chromosome conformation capture by Hi-C

The Hi-C protocol was performed as described by[82]. 10 ml of Bj109 cell culture at exponential phase in AG medium, was passed through a 0.5-mm needle several times to filter out excess exopolysaccharide, and then this cell suspension was used for subsequent steps. Protein-DNA interactions were cross-linked by adding formaldehyde (Sigma-Aldrich) to a final concentration of 3% and incubating for 30 minutes at room temperature (RT), with gentle agitation. Crosslinking was quenched with glycine (250 mM final concentration) for 20 min at RT with gentle agitation. After two washes with 1x PBS, pellets were stored at −70°C until further use. Frozen pellets were thawed on ice and resuspended in 1mL of TE buffer (Sigma-Aldrich, Cat# T928) containing a protease inhibitor tablet (Roche, Cat# 11873580001). Subsequently, 12 µL of Ready-Lyse lysozyme (LGC Biosearch, Cat# LU-R1804M) and 500 U of mutanolysin (M4748-5KU, Sigma-Aldrich) were added, and this mixture was incubated at 37°C for 45 min with gentle agitation. SDS was added to a final concentration of 0.5% and the samples were incubated for an additional 10 min at RT with gentle agitation. Samples were prepared for digestion by adding 3000µL water, 500 µL NEB3 10× digestion buffer, and 500 µL Triton X-100 (10%). A 400 µL non-digested control was aliquoted, and 1000 U of HpaII was added to the sample, followed by incubation for 3 h at 37°C with agitation (180 rpm, INFORS Unitron standard). After incubation, 400 µL of the sample was reserved as a digestion control, and the remaining sample was centrifuged at 4°C and resuspended in 400 µL of water.

Biotinylation was carried out for 1 h at 37°C with agitation (180 rpm, INFORS Unitron standard) after the addition of a mix of 50 µL 10x Ligation Buffer (500 mM Tris-HCl pH 7.5, 100 mM MgCl2, 100 mM DTT), 4.5 µL 10 mM dAGTTp, 37.5 µL Biotin-14-dCTP (Thermo Fisher) and 8 µL Large (Klenow) Fragment DNA Polymerase (5 U/µL, NEB). Chromatin ligation was performed with a mix of 120 µL Thermo Scientific T4 10× ligation buffer, 12 µL BSA (10 mg/mL), 12 µL ATP (100 mM), 16 µL T4 DNA Ligase (30 U/µL), and 540 µL of water. The mixture was incubated for 3 h at RT with gentle agitation, allowing ligation. Following ligation, protein-DNA cross-links were reversed in the samples and digested and non-digested controls by the addition of Proteinase K, EDTA (6 mM final concentration) and SDS (0.5% final concentration) followed by an overnight incubation at 65°C. DNA was purified with phenol:chloroform:isoamyl alcohol (25:24:1). Libraries were constructed as described in[83], including DNA shearing (Covaris S220), end-repair/dA-tailing, adapter ligation, and library amplification. Libraries were then pooled in equimolar proportions and sequenced as 150 bp paired-end reads on Illumina NovaSeq X Plus Series.

For Hi-C analysis, contact maps were generated using the OCHA pipeline in R[84], using the complete genome sequence of BjE109, available at NCBI GenBank under the accession no. CP010313[54].The remaining data were binned into sparse contact matrices and normalized using the balanced normalization method. The Hi-C map shown in Figure 5 is the result of the merge of 4 Hi-C libraries which 3 of them were biological replicates.

## Data availability

Deep sequencing data from Hi-C were submitted to the GenBank Sequence Read Archive (SRA) as BioProject PRJNA1557419.

## Author contributions

**Conceptualization:** VSL & ASB; **Data Curation:** VSL, ASB, IM; **Formal Analysis:** IM, ASB, JB, EJM, VSL; **Funding acquisition:** ASB, DJC, VSL; **Investigation:** IM, JB, EJM, ASB, VSL; **Methodology:** IM, ASB, EJM, VSL; **Supervision:** ASB, EJM, DJC, VSL; **Writing**: IM, ASB, VSL.

## Acknowledgments

We want to thank the several programs allowing us to exchange between laboratories: Interdisciplinary Center MICROBES of the Université Paris-Saclay, as part of the France 2030 program ANR-11-IDEX-0003 (support of MICROBES mobility program in 2023 and 2025 (to IM), and Missionaires Invités 2025 (to ASB) from the Université Paris-Saclay; Programa de Movilidad Internacional – Universidad Nacional de San Martín (to IM); Centro Franco Argentino-Universidad Nacional de San Martín Ambassade de France in Argentine and Institut Francais Argentine (to VSL). This study was supported by the International Center for Genetic Engineering and Biotechnology (CRP/ARG18-06_EC to A.S.B.), the Agencia Nacional de Promoción de la Investigación, el Desarrollo Tecnológico y la Innovación of Argentina (PICT-2020-SERIEA-00521 and PICT-2021-CatI-0154 Aplicación intensivo, PIP 2022-2024-CONICET to A.S.B.) and UNSAM (UNSAM Investiga 2025-Subsidio 12). Additional support came from the Interdisciplinary Center MICROBES of the Université Paris-Saclay, as part of the France 2030 program ANR-11-IDEX-0003 (to V.S.L, program Deep-HiC). A.S.B. and D.J.C. are Career Members of CONICET. IM and J.B. are CONICET scholarship recipients. We are grateful to Leticia Larotonda, Nicolás Ríos, Soledad Guidolín, Inés Marchesini, Frédéric Boccard and Stéphanie Bury-Moné for useful discussions. We thank Mohamed Seba, Elisa Galli and Sokrich Ponndara for their technical help. We also want to thank Pascaline Tirand and Fanny Culot for daily help and preparing culture media. We acknowledge the use of AI to improve the English in this manuscript.

## Reference

1. Jun, S., et al., Fundamental principles in bacterial physiology-history, recent progress, and the future with focus on cell size control: a review. Rep Prog Phys, 2018. 81(5): p. 056601.

2. Monod, J., The Growth of Bacterial Cultures. Annual Reviews in Microbiology, 1949. 3: p. 371–394.

3. Lane-Claypon, J.E., Multiplication of Bacteria and the Influence of Temperature and some other Conditions thereon. J Hyg (Lond), 1909. 9(2): p. 239–48.

4. Govers, S.K. and C. Jacobs-Wagner, Caulobacter crescentus: model system extraordinaire. Curr Biol, 2020. 30(19): p. R1151–R1158.

5. Willis, L. and K.C. Huang, Sizing up the bacterial cell cycle. Nat Rev Microbiol, 2017. 15(10): p. 606–620.

6. Madin, J.S., et al., A synthesis of bacterial and archaeal phenotypic trait data. Sci Data, 2020. 7(1): p. 170.

7. Kysela, D.T., et al., Diversity Takes Shape: Understanding the Mechanistic and Adaptive Basis of Bacterial Morphology. PLoS Biol, 2016. 14(10): p. e1002565.

8. Hoffert, M., et al., Constructing a “periodic table” of bacteria to map diversity in trait space. ISME J, 2026. 20(1).

9. Aliperti, L., et al., r/K selection of GC content in prokaryotes. Environ Microbiol, 2023. 25(12): p. 3255–3268.

10. Couso, L.L., et al., Ecology theory disentangles microbial dichotomies. Environ Microbiol, 2023. 25(12): p. 3052–3063.

11. Soler-Bistue, A., L.L. Couso, and I.E. Sanchez, The evolving copiotrophic/oligotrophic dichotomy: From Winogradsky to physiology and genomics. Environ Microbiol, 2023.

12. Wang, P., et al., Robust growth of Escherichia coli. Curr Biol, 2010. 20(12): p. 1099–103.

13. Elf, J. and I. Barkefors, Single-Molecule Kinetics in Living Cells. Annu Rev Biochem, 2019. 88: p. 635–659.

14. Wallden, M., et al., The Synchronization of Replication and Division Cycles in Individual E. coli Cells. Cell, 2016. 166(3): p. 729–739.

15. Henson, M.W., et al., Cultivation and genomics of the first freshwater SAR11 (LD12) isolate. ISME J, 2018. 12(7): p. 1846–1860.

16. Le, T.B., et al., High-resolution mapping of the spatial organization of a bacterial chromosome. Science, 2013. 342(6159): p. 731–4.

17. Marbouty, M., et al., Condensin- and Replication-Mediated Bacterial Chromosome Folding and Origin Condensation Revealed by Hi-C and Super-resolution Imaging. Mol Cell, 2015. 59(4): p. 588–602.

18. Lioy, V.S., et al., Multiscale Structuring of the E. coli Chromosome by Nucleoid-Associated and Condensin Proteins. Cell, 2018. 172(4): p. 771–783 e18.

19. Goley, E.D., et al., Dynamic chromosome organization and protein localization coordinate the regulatory circuitry that drives the bacterial cell cycle. Cold Spring Harb Symp Quant Biol, 2009. 74: p. 55–64.

20. Pawlikiewicz, K., et al., SMC ensures efficient chromosome replication and oriC positioning during Streptomyces spore germination. Sci Rep, 2026. 16(1).

21. Plaskowska, K. and J. Zakrzewska-Czerwinska, Chromosome structure and DNA replication dynamics during the life cycle of the predatory bacterium Bdellovibrio bacteriovorus. FEMS Microbiol Rev, 2023. 47(6).

22. Jalal, A.S.B. and T.B.K. Le, Bacterial chromosome segregation by the ParABS system. Open Biol, 2020. 10(6): p. 200097.

23. Fogel, M.A. and M.K. Waldor, A dynamic, mitotic-like mechanism for bacterial chromosome segregation. Genes Dev, 2006. 20(23): p. 3269–82.

24. Toro, E., et al., Caulobacter requires a dedicated mechanism to initiate chromosome segregation. Proc Natl Acad Sci U S A, 2008. 105(40): p. 15435–40.

25. Ptacin, J.L., et al., A spindle-like apparatus guides bacterial chromosome segregation. Nat Cell Biol, 2010. 12(8): p. 791–8.

26. Lim, H.C., et al., Evidence for a DNA-relay mechanism in ParABS-mediated chromosome segregation. Elife, 2014. 3: p. e02758.

27. Livny, J., Y. Yamaichi, and M.K. Waldor, Distribution of centromere-like parS sites in bacteria: insights from comparative genomics. J Bacteriol, 2007. 189(23): p. 8693–703.

28. Wang, X., et al., Bacillus subtilis SMC complexes juxtapose chromosome arms as they travel from origin to terminus. Science, 2017. 355(6324): p. 524–527.

29. Wang, X., et al., Condensin promotes the juxtaposition of DNA flanking its loading site in Bacillus subtilis. Genes Dev, 2015. 29(15): p. 1661–75.

30. Vallet-Gely, I. and F. Boccard, Chromosomal organization and segregation in Pseudomonas aeruginosa. PLoS Genet, 2013. 9(5): p. e1003492.

31. Lioy, V.S., et al., Distinct Activities of Bacterial Condensins for Chromosome Management in Pseudomonas aeruginosa. Cell Rep, 2020. 33(5): p. 108344.

32. Sanath-Kumar, R., et al., Structural Maintenance of Chromosomes Complexes in Bacteria. Annu Rev Microbiol, 2026. 80(1): p. 243–262.

33. Sullivan, N.L., K.A. Marquis, and D.Z. Rudner, Recruitment of SMC by ParB-parS organizes the origin region and promotes efficient chromosome segregation. Cell, 2009. 137(4): p. 697–707.

34. Gruber, S. and J. Errington, Recruitment of condensin to replication origin regions by ParB/SpoOJ promotes chromosome segregation in B. subtilis. Cell, 2009. 137(4): p. 685–96.

35. Delgado-Baquerizo, M., et al., A global atlas of the dominant bacteria found in soil. Science, 2018. 359(6373): p. 320–325.

36. Hungria, M. and I.C. Mendes, Nitrogen Fixation with Soybean: The Perfect Symbiosis?, in Biological Nitrogen Fixation. 2015, John Wiley & Sons, Inc. p. 1009–1024.

37. Santos, M.S., M.A. Nogueira, and M. Hungria, Microbial inoculants: reviewing the past, discussing the present and previewing an outstanding future for the use of beneficial bacteria in agriculture. AMB Express, 2019. 9(1): p. 205.

38. Sethu Madhavan, A. and L.M. Muller, Balancing mutualism: choice and sanctions in root-microbe symbioses. New Phytol, 2026.

39. Marcondes de Souza, J.A., et al., The Family Bradyrhizobiaceae, in The Prokaryotes Alphaproteobacteria and Betaproteobacteria, E. Rosenberg, et al., Editors. 2014, Springer Nature: Springer Berlin, Heidelberg. p. 135–154.

40. Medici, I.F., et al., The distinct cell physiology of Bradyrhizobium at the population and cellular level. BMC Microbiol, 2024. 24(1): p. 129.

41. Sarao, S.K., et al., Bradyrhizobium diazoefficiens cultures display phenotypic heterogeneity. ISME Commun, 2025. 5(1): p. ycaf054.

42. Tomasch, J., S. Koppenhofer, and A.S. Lang, Connection Between Chromosomal Location and Function of CtrA Phosphorelay Genes in Alphaproteobacteria. Front Microbiol, 2021. 12: p. 662907.

43. Amstutz, J., et al., Getting to the point: unipolar growth of Hyphomicrobiales. Curr Opin Microbiol, 2024. 79: p. 102470.

44. Williams, M.A., D.T. Kysela, and P.J. Brown, Diversity of growth patterns in the Alphaproteobacteria, in Cell cycle regulation and development in alphaproteobacteria, E.G. Biondi, Editor. 2022, Springer International Publishing: Cham: Springer International Publishing. p. pp. 185-220.

45. Ostermayer, J., et al., Taming wild replicons: evolution and domestication of large extrachromosomal replicons. Curr Opin Microbiol, 2025. 88: p. 102657.

46. Czarnecki, J., et al., Large extrachromosomal replicons are widespread across bacterial lineages and show coordinated replication termination and spatial coupling with the chromosome. Nat Commun, 2026.

47. Hsu, Y.P., et al., Full color palette of fluorescent d-amino acids for in situ labeling of bacterial cell walls. Chem Sci, 2017. 8(9): p. 6313–6321.

48. Kuru, E., et al., Synthesis of fluorescent D-amino acids and their use for probing peptidoglycan synthesis and bacterial growth in situ. Nat Protoc, 2015. 10(1): p. 33–52.

49. Evertts, A.G. and H.A. Coller, Back to the origin: reconsidering replication, transcription, epigenetics, and cell cycle control. Genes Cancer, 2012. 3(11-12): p. 678–96.

50. Hutchison, C.A., 3rd, et al., Design and synthesis of a minimal bacterial genome. Science, 2016. 351(6280): p. aad6253.

51. de Lorenzo, V., It’s the metabolism, stupid! Environ Microbiol Rep, 2015. 7(1): p. 18–9.

52. Liu, X., J. Yan, and M.W. Kirschner, Cell size homeostasis is tightly controlled throughout the cell cycle. PLoS Biol, 2024. 22(1): p. e3002453.

53. Sanders, S., et al., Beyond the average: An updated framework for understanding the relationship between cell growth, DNA replication, and division in a bacterial system. PLoS Genet, 2023. 19(1): p. e1010505.

54. Fu, J., et al., Cells prioritize the regulation of cell mass density. Sci Adv, 2025. 11(35): p. eadv9759.

55. Cezanne, A., et al., The Archaeal Cell Cycle. Annu Rev Cell Dev Biol, 2024. 40(1): p. 1–23.

56. Szafran, M.J., et al., Spatial rearrangement of the Streptomyces venezuelae linear chromosome during sporogenic development. Nat Commun, 2021. 12(1): p. 5222.

57. Weber, P.M., et al., A Bidimensional Segregation Mode Maintains Symbiont Chromosome Orientation toward Its Host. Curr Biol, 2019. 29(18): p. 3018–3028 e4.

58. Niault, T., et al., Cell cycle-coordinated maintenance of the Vibrio bipartite genome. EcoSal Plus, 2023. 11(1): p. eesp00082022.

59. Singhi, D., et al., Rhodoccoccus erythropolis Is Different from Other Members of Actinobacteria: Monoploidy, Overlapping Replication Cycle, and Unique Segregation Pattern. J Bacteriol, 2019. 201(24).

60. Trojanowski, D., et al., Choreography of the Mycobacterium replication machinery during the cell cycle. mBio, 2015. 6(1): p. e02125–14.

61. Fournes, F., et al., Genome replication and cell division control by the methylation-sensitive GcrA regulator in Agrobacterium tumefaciens. BioRxiv, 2026. 728657v1.

62. Ehrle, H.M., et al., Polar Organizing Protein PopZ Is Required for Chromosome Segregation in Agrobacterium tumefaciens. J Bacteriol, 2017. 199(17).

63. Wagner, M., et al., Engineering a Sinorhizobium meliloti Chassis with Monopartite, Single Replicon Genome Configuration. ACS Synth Biol, 2024. 13(8): p. 2515–2532.

64. Krol, E., et al., Stable inheritance of Sinorhizobium meliloti cell growth polarity requires an FtsN-like protein and an amidase. Nat Commun, 2021. 12(1): p. 545.

65. Deghelt, M., et al., G1-arrested newborn cells are the predominant infectious form of the pathogen Brucella abortus. Nat Commun, 2014. 5: p. 4366.

66. Francis, N., et al., CtrA controls cell division and outer membrane composition of the pathogen Brucella abortus. Mol Microbiol, 2017. 103(5): p. 780–797.

67. Torres, D., et al., Genome Sequence of Bradyrhizobium japonicum E109, One of the Most Agronomically Used Nitrogen-Fixing Rhizobacteria in Argentina. Genome Announc, 2015. 3(1).

68. Brown, P.J., et al., Polar growth in the Alphaproteobacterial order Rhizobiales. Proc Natl Acad Sci U S A, 2012. 109(5): p. 1697–701.

69. Ren, Z., et al., Centromere Interactions Promote the Maintenance of the Multipartite Genome in Agrobacterium tumefaciens. mBio, 2022. 13(3): p. e0050822.

70. Frage, B., et al., Spatiotemporal choreography of chromosome and megaplasmids in the Sinorhizobium meliloti cell cycle. Mol Microbiol, 2016. 100(5): p. 808–23.

71. Izquierdo-Martinez, A., et al., Chromosome segregation dynamics during the cell cycle of Staphylococcus aureus. Nat Commun, 2025. 16(1): p. 8626.

72. Brilli, M., et al., The diversity and evolution of cell cycle regulation in alpha-proteobacteria: a comparative genomic analysis. BMC Syst Biol, 2010. 4: p. 52.

73. Pini, F., et al., The DivJ, CbrA and PleC system controls DivK phosphorylation and symbiosis in Sinorhizobium meliloti. Mol Microbiol, 2013. 90(1): p. 54–71.

74. Pini, F., et al., Cell Cycle Control by the Master Regulator CtrA in Sinorhizobium meliloti. PLoS Genet, 2015. 11(5): p. e1005232.

75. Dendene, S., et al., Sinorhizobium meliloti FcrX coordinates cell cycle and division during free-living growth and symbiosis by a ClpXP-dependent mechanism. Proc Natl Acad Sci U S A, 2025. 122(11): p. e2412367122.

76. Martin, S., et al., DNA methylation by CcrM contributes to genome maintenance in the Agrobacterium tumefaciens plant pathogen. Nucleic Acids Res, 2024. 52(19): p. 11519–11535.

77. Lamouche, F., et al., From Intracellular Bacteria to Differentiated Bacteroids: Transcriptome and Metabolome Analysis in Aeschynomene Nodules Using the Bradyrhizobium sp. Strain ORS285 bclA Mutant. J Bacteriol, 2019. 201(17).

78. Chen, W.F., et al., Bacteroid Development, Transcriptome, and Symbiotic Nitrogen-Fixing Comparison of Bradyrhizobium arachidis in Nodules of Peanut (Arachis hypogaea) and Medicinal Legume Sophora flavescens. Microbiol Spectr, 2023. 11(1): p. e0107922.

79. Greetatorn, T., et al., Differential responses of Bradyrhizobium sp. SUTN9-2 to plant extracts and implications for endophytic interactions within different host plants. Sci Rep, 2025. 15(1): p. 3154.

80. Quelas, J.I., et al., Analysis of two polyhydroxyalkanoate synthases in Bradyrhizobium japonicum USDA 110. J Bacteriol, 2013. 195(14): p. 3145–55.

81. Ducret, A., E.M. Quardokus, and Y.V. Brun, MicrobeJ, a tool for high throughput bacterial cell detection and quantitative analysis. Nat Microbiol, 2016. 1(7): p. 16077.

82. Cockram, C., et al., Euryarchaeal genomes are folded into SMC-dependent loops and domains, but lack transcription-mediated compartmentalization. Mol Cell, 2021. 81(3): p. 459–472 e10.

83. Lioy, V.S., et al., Dynamics of the compartmentalized Streptomyces chromosome during metabolic differentiation. Nat Commun, 2021. 12(1): p. 5221.

84. Serizay, J., et al., Orchestrating chromosome conformation capture analysis with Bioconductor. Nat Commun, 2024. 15(1): p. 1072.

