## Supplementary Material for "Asymmetric division and polar growth define a non-canonical cell cycle in *Bradyrhizobium*"

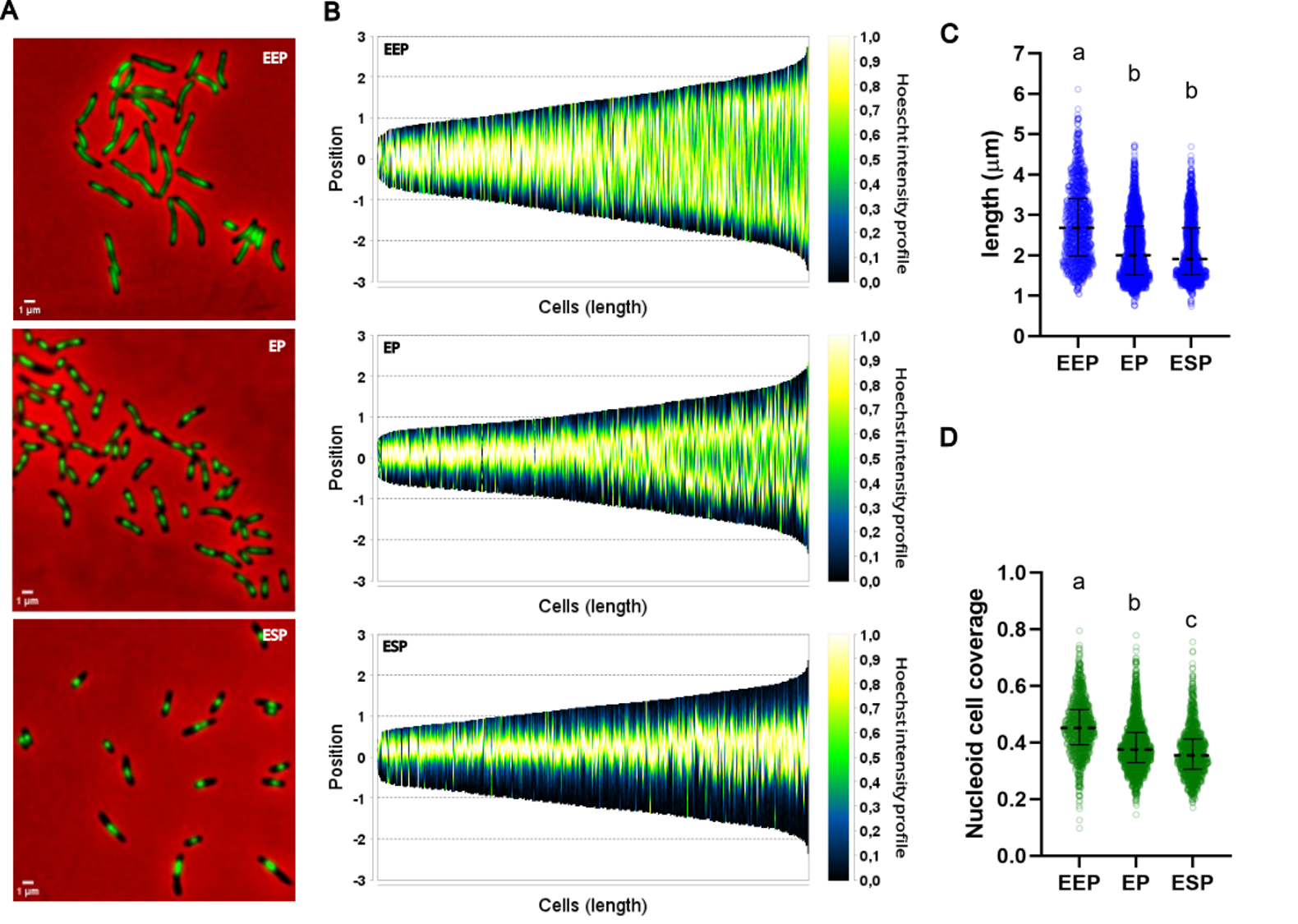

**Figure S1. Cell size and nucleoid distribution vary along the growth phases of Bj109.** **A.** Representative image of cells stained with Hoechst 33342 at different stages of growth in AG medium. **B.** Demographic representation showing Hoechst 33342 intensity profile of cells as a function of cell length at the indicated phases. **C.** Distribution of cell sizes at the indicated phases. **D.** Proportion of the cell covered by Hoechst 33342 intensity at the indicated phases. The middle lines indicate the median for each timepoint. Bars show the quartile range. Letters indicate statistically different groups (Kruskal Wallis and Dunn’s test, p<0,0001). For panels B to D the number of analysed cells is 1309 for EEP, 1717 for EP and 1444 for ESP.

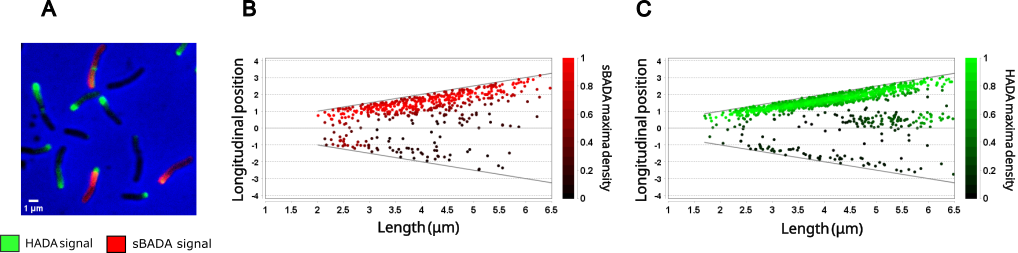

**Figure S2. Cell wall synthesis shows a sequential pattern. A.** Representative fluorescence microscopy image of Bj109 cells during exponential phase in AG medium. Cells were sequentially stained with the FDAAs sBADA and then HADA. **B & C.** Distribution and density of sBADA and HADA maxima, within cells sorted by length and oriented by maximum fluorescence intensity of each channel (N = 1395).

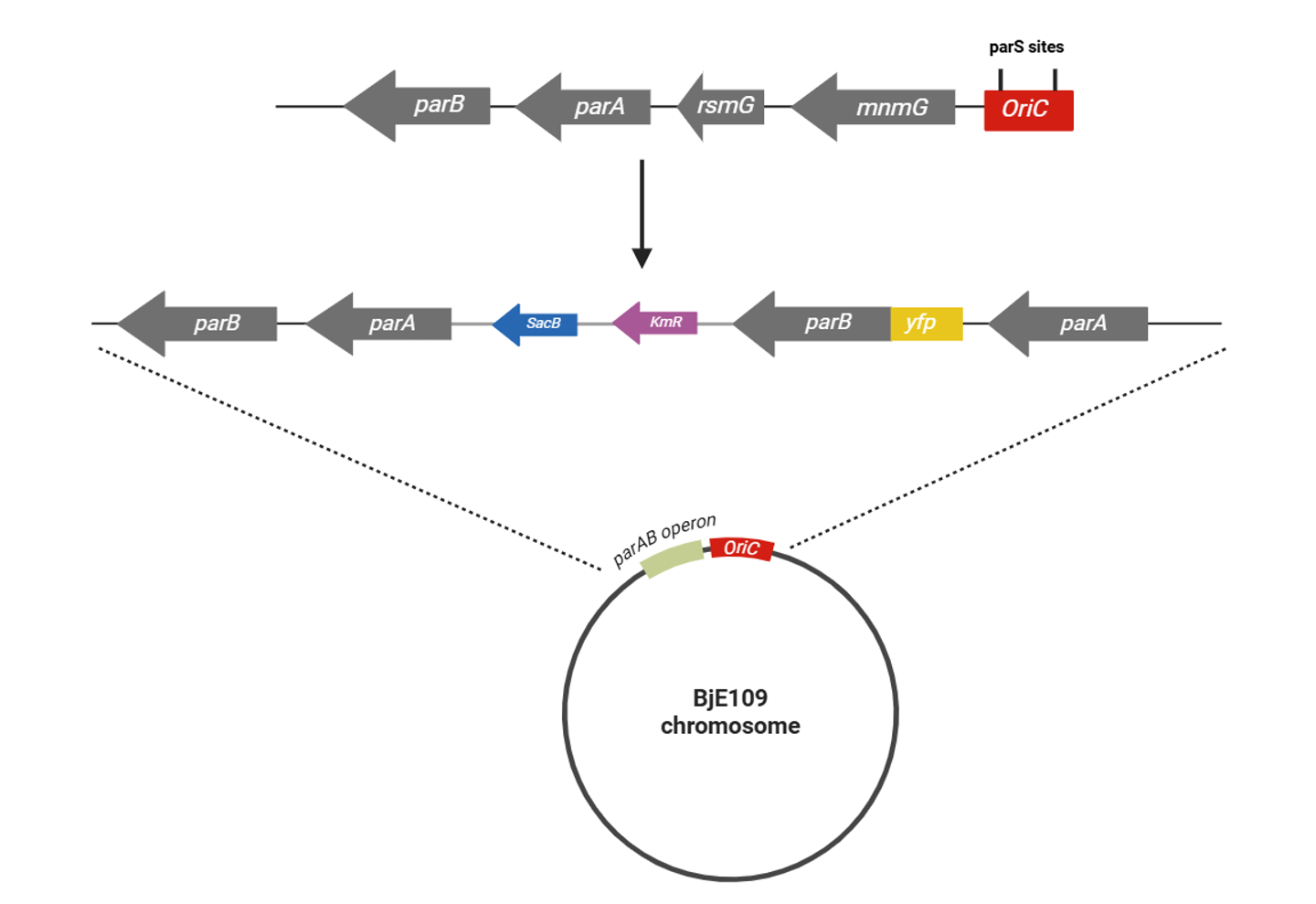

**Figure S3.** **Schematic representation of the genomic modifications incorporated in the *yfp-parB* merodiploid strain.** This strain contains an *yfp-parB* fusion and an additional copy of *parA* at the parABS locus, that was inserted by recombination of the vector pK18mobSacB containing the *parA/yfp-parB* construction.

**Supplementary text:**

***parAB* merodiploid construction:** *parB* in Bj109 chromosome corresponds to locus tag RN69_02930 in the complementary strand between 619,483-620,370, close to the *oriC* (located at 624,080 to 624,665 nt , DOI:[10.1093/nar/gkac964](https://doi.org/10.1093/nar/gkac964)), with a predicted 295 amino acids product with high similarity to *Caulobacter, Brucella* and *Sinorhizobium* alleles. It is also located downstream the *parA* gene.

Meanwhile, there is a second putative copy of the gene in the locus tag RN69_29295. However, this gene is located far from the *oriC* (6,144,299-6,144,928, corresponding to 2/3 of the left replichore), resembling a “ParB-like protein” but encoding for a shorter peptide of 208 amino acids, that while it shows similarity with other parB-like genes from *Caulobacter* it does not present orthologs encoded in *Brucella* or *Sinorhizobium*. Also, the gene is not physically associated with a *parA* gene but rather isolated.

Therefore, we constructed a YFP-ParB translational fusion using Gibson Assembly Master Mix (NEB) and cloned into pK18*mobsacB*, following the manufacturer’s protocols. For this, we amplified sYFP2 with adapted codons for Bj109, from pRJPaph-sYFP2 (Addgene plasmid # 67033). The *parA* coding sequence including *parB* upstream region and *parB* coding sequence were amplified from Bj109 genomic DNA. The primers for amplifying these DNA fragments were designed following instructions from NEBuilder tool, to be able to insert them in the linearized pK18*mobsacB* cut with *EcoRI* and *PstI*. The final construction, pK18*mobsacB-parA/yfp-parB*, was transformed into *E. coli* s17-1 λ*pir* to finally introduce it in BjE109 by conjugation.

Since in *Brucella abortus* this fusion is deleterious (DOI: 10.1038/ncomms5366), we directly chose to insert the *yfp-parB* gene into the chromosome in tandem downstream the *parAB* locus (Figure S3). Also, we added a second *parA* copy to avoid altering ParA:ParB stoichiometry.

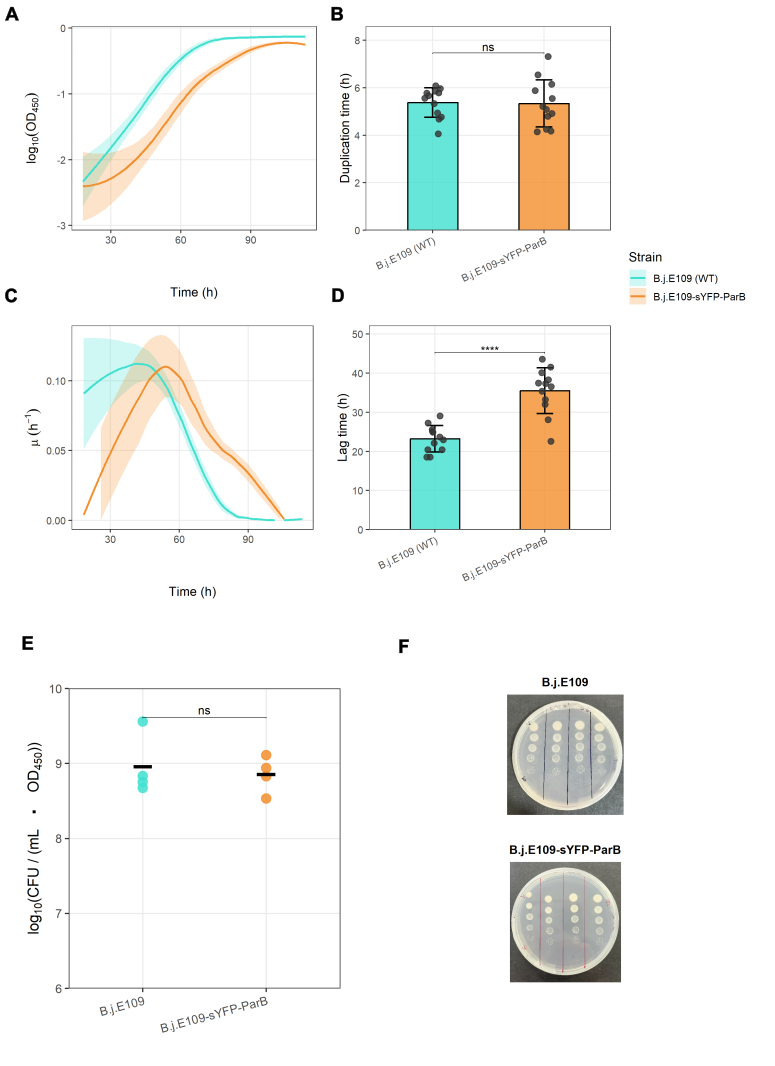

**Figure S4. Bj109-YFPParB strain presents similar physiological parameters to wild type strain. A.** Growth curve of wild type strain (green) and YFP-ParB derivative (orange). **B.** Generation time of both strains calculated from two independent experiments with at list six technical replicates each, of automatic growth curves. **C.** Growth rate as a function of time of growth curves. **D.** Lag time of both strains calculated from two independent experiments with at list six technical replicates each, of automatic growth curves. **E.** Viability test of both strains showing a non-statistical difference in CFU count per ml and OD unit. **F.** Representative plate of the experiment shown in E.

**Figure S5:** **ParB is located at the boundaries of DNA regions towards the poles of the cell.** **A.** Representative fluorescence microscopy images of cells grown in AG medium at exponential phase. Cells expressing YFP-ParB were stained with Hoesch for DNA visualization. **B.** Demographic representation showing the YFP-ParB and Hoesch intensity profile of cells sorted by length and oriented by nucleoid position (N = 2867). Distribution of nucleoid centers **(C)** and YFP-ParB foci **(D)** within the cells sorted by length. **E.** Cell cycle and chromosome segregation representation schemed according to the analyses of microscopy images.
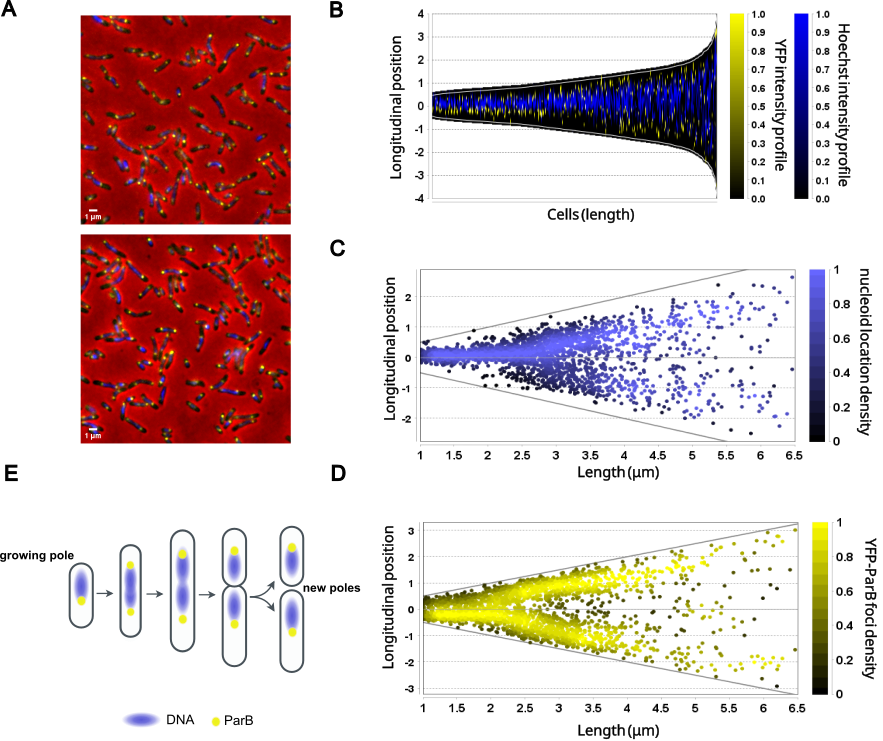

**Table S1:** Bacterial strains used in this study.

| **Strain** | **Relevant characteristics** | **Source** |
| --- | --- | --- |
| *B. japonicum* E109 | *B. japonicum* E109 wild type | Fabricio Cassán Laboratory |
| *B. japonicum* E109 – parA/yfp-parB | *B. japonicum* E109 expressing translational fusion of YFP-ParB | This study |
| *E. coli* s17-1 λpir | Used for transformation by biparental conjugation. | Laboratory stock |
| *E. coli* DH5α | Used for plasmid preparation | Laboratory stock |

**TableS2**. Descriptive statistics of cell length, nucleoid coverage and nucleoid positional asymmetry in AG medium from early exponential (EEP), exponential (EP) and early stationary phases (ESP). To calculate the distance of nucleoids to midcell, the feature recognition tool of MicrobeJ was used, and the location relative to the longitudinal axes was obtained from cells bearing only one nucleoid.

| **Cell length analysis** | **EEP** | **EP** | **ESP** |
| --- | --- | --- | --- |
| N | 547 | 759 | 861 |
| Minimum | 1.015 | 1.012 | 1.062 |
| 25% Percentile | 2.011 | 1.698 | 1.685 |
| Median | 2.784 | 2.091 | 2.312 |
| 75% Percentile | 3.428 | 2.463 | 3.014 |
| Maximum | 6.533 | 5.298 | 5.564 |
| Mean | 2.795 | 2.144 | 2.440 |
| Std. Deviation | 0.948 | 0.602 | 0.905 |
| Std. Error of Mean | 0.041 | 0.022 | 0.031 |
| **Nucleoid coverage** | **EEP** | **EP** | **ESP** |
| N | 528 | 759 | 861 |
| Minimum | 0.0194 | 0.0754 | 0.0277 |
| 25% Percentile | 0.4179 | 0.2956 | 0.2533 |
| Median | 0.5099 | 0.3749 | 0.3225 |
| 75% Percentile | 0.5961 | 0.4709 | 0.3924 |
| Maximum | 0.8791 | 0.8494 | 0.9265 |
| Range | 0.8597 | 0.7740 | 0.8988 |
| Mean | 0.5084 | 0.3860 | 0.3263 |
| Std. Deviation | 0.1256 | 0.1289 | 0.1254 |
| Std. Error of Mean | 0.0055 | 0.0047 | 0.0043 |
| **Relative distance to midcell of nucleoid position** | **EEP** | **EP** | **ESP** |
| Mean +/- SD | 0.11±0.1 | 0.11±0.08 | 0.22±0.14 |

**TableS3**. Descriptive statistics of cell length, nucleoid coverage and nucleoid positional asymmetry in YEM media from (EEP), exponential (EP) and early stationary phases (ESP). To calculate the distance of nucleoids to midcell, the feature recognition tool of MicrobeJ was used, and the location relative to the longitudinal axes was obtained from cells bearing only one nucleoid.

| **Cell length analysis** | **EEP** | **EP** | **ESP** |
| --- | --- | --- | --- |
| N | 1309 | 1978 | 1444 |
| Minimum | 1.033 | 0.751 | 0.739 |
| 25% Percentile | 1.988 | 1.519 | 1.527 |
| Median | 2.678 | 2.002 | 1.912 |
| 75% Percentile | 3.411 | 2.719 | 2.683 |
| Maximum | 6.113 | 4.734 | 4.691 |
| Mean | 2,779 | 2,168 | 2,143 |
| Std. Deviation | 0,9334 | 0,7679 | 0,7509 |
| Std. Error of Mean | 0,02580 | 0,01727 | 0,01976 |
| **Nucleoid coverage analysis** | **EEP** | **EP** | **ESP** |
| N | 1309 | 1862 | 1276 |
| Minimum | 0.09706 | 0.1457 | 0.1699 |
| 25% Percentile | 0.3919 | 0.3287 | 0.3051 |
| Median | 0.4510 | 0.3758 | 0.3547 |
| 75% Percentile | 0.5166 | 0.4355 | 0.4123 |
| Maximum | 0.7947 | 0.7785 | 0.7551 |
| Range | 0.6976 | 0.6327 | 0.5852 |
| Mean | 0.4556 | 0.3889 | 0.3652 |
| Std. Deviation | 0.09368 | 0.08873 | 0.08548 |
| Std. Error of Mean | 0.002589 | 0.002056 | 0.002393 |
| **Relative distance to midcell of nucleoid position** | **EEP** | **EP** | **ESP** |
| Mean +/- SD | 0.07±0.16 | 0.13±0.09 | 0.21±0.12 |
